# Songbird ventral basal ganglia sends performance error signals to dopaminergic midbrain

**DOI:** 10.1101/346841

**Authors:** Ruidong Chen, Pavel A. Puzerey, Andrea C. Roeser, Tori E. Riccelli, Archana Podury, Kamal Maher, Alexander Farhang, Jesse H. Goldberg

## Abstract

Ventral tegmental area (VTA) dopamine neurons signal prediction error, the difference between actual and predicted outcome, but it remains unclear how error is computed. Here we identify in songbirds a ventral basal ganglia (vBG) region that is required for song learning and that sends prediction error signals to VTA. During singing, vBG neurons heterogeneously encoded song timing, auditory error, predicted error, and the difference between the two (prediction error). Viral tracing revealed inputs to vBG from auditory and vocal motor thalamus, auditory and vocal motor cortex, and VTA. Our findings reveal a classic actor-critic circuit motif in which a ventral critic learns the ‘prediction’ component of a prediction error signal that is relayed by VTA to a dorsal actor (the vocal motor BG nucleus Area X). A circuit motif for computing reward prediction error can compute predicted performance quality during motor sequence learning.

## INTRODUCTION

When practicing motor performance such as a piano concerto, you may aspire to sound like Glenn Gould playing Chopin’s Prelude No. 4. But one problem is that your performance will reliably fall short of your aspirations. Trial and error learning proceeds via incremental improvement in performance quality, requiring performance to be evaluated against benchmarks that change with practice (Schmidt et al., 2018; Thelen, 1995). Neural mechanisms of performance benchmarking and evaluation remain poorly understood.

Reinforcement theory may offer insights into performance evaluation during motor sequence learning (Sutton and Barto, 1998). In classic actor-critic (AC) models (Figure 1A), an ‘actor’ module learns to select the reward-maximizing action in a given ‘state’, where state can be a spatial location, a sensory cue, or elapsed time from cue onset (Joel et al., 2002; Suri and Schultz, 1998). The actor’s learning rule is to modify the connections between state and action choices according to a reinforcement signal from a separate ‘critic’. The job of the critic is to generate this reinforcement signal, often in the form of reward prediction error (RPE), the difference between the reward received and the reward predicted based on the state. To signal RPE, the critic learns a state-dependent reward prediction and compares this prediction to the actual reward received.

**Figure 1.**
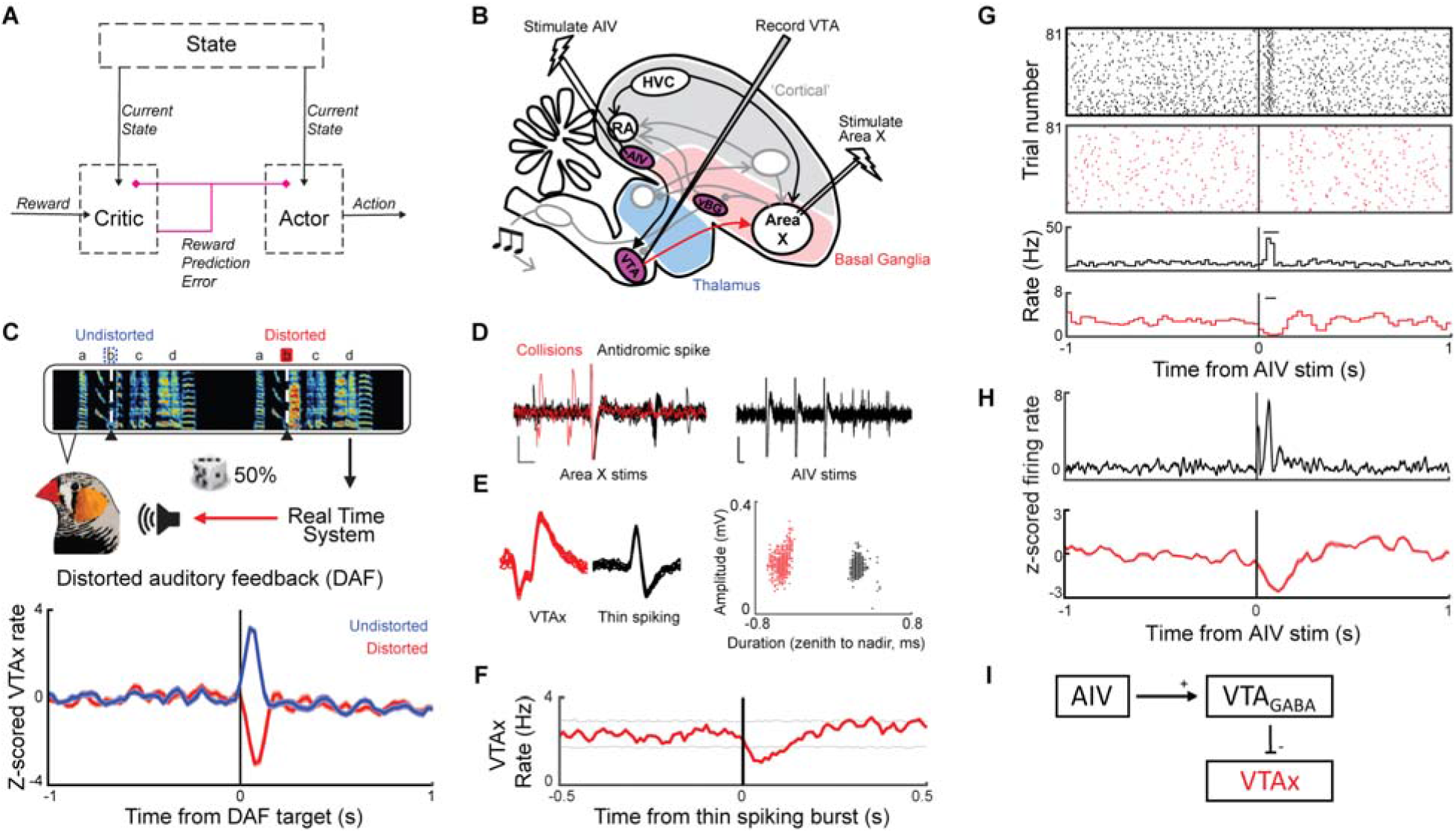
An auditory cortical projection suppresses VTAx neurons through feedforward inhibition. (A) Schematic of the Actor-Critic network. (B) Syllable-targeted DAF (top) results in dopaminergic prediction error signals (bottom, from ref (Gadagkar et al., 2016)). (C) Test of VTAx response to AIV stimulation. (D to G) Experiment conducted on simultaneously recorded wide-spiking VTAx neuron and thin-spiking neuron. (D) Antidromic identification (left), and AIV stimulation (right). (E) Units in (D-G) exhibited distinct waveforms. (F) Cross-correlogram of spontaneous firing between the two units. Dotted lines, 95% confidence interval (STAR Methods). (G) Raster plots (top) and rate histograms (bottom) of thin spiking (black) and VTAx neuron (red), aligned to AIV stimulation. Horizontal bars indicate significant response (p<0.05, Z test) (STAR Methods). (H) Average Z-scored response to AIV stimulation from 4 VTAx (red) and 7 thin-spiking neurons (black). (I) Summary: AIV can inhibit VTAx neurons by activating local interneurons.

The discovery of RPE signaling by VTA dopamine (DA) neurons suggested that AC-like networks may be biologically realized in mammalian brains (Montague et al., 2004; Schultz et al., 1997). VTA DA signals reach both dorsal and ventral divisions of the mammalian BG (Haber et al., 2000), which may instantiate parts of the actor and critic, respectively (Atallah et al., 2007; O’Doherty et al., 2004; Takahashi et al., 2008). Consistent with AC model predictions, both ventral and dorsal BG receive information about current state from cortical inputs, receive RPE reinforcement signals from dopaminergic midbrain, and can learn value-weighted state representations by implementing DA-modulated corticostriatal plasticity (Bar-Gad et al., 2000; Haber et al., 2000; Houk and Wise, 1995). The ventral basal ganglia (vBG) is thought to play a part of the critic because it can signal state-dependent reward prediction to downstream VTA neurons, providing the critic’s output with the ‘prediction’ component of reward prediction error (Takahashi et al., 2016). The dorsal BG is thought to play the actor because it can signal action value and bias downstream motor circuits to generate reward maximizing actions (Hikosaka et al., 2006; Lauwereyns et al., 2002; Samejima et al., 2005; Tai et al., 2012). These studies in mammals have focused on reward seeking, and it remains unknown if similar principles would apply to motor learning problems, such as speech or music, where performance outcomes are compared to internal benchmarks, and not to reward.

Songbirds provide a tractable model system to identify mechanisms of performance evaluation and learning. Zebra finches learn to imitate a sequence of song notes, or syllables, acquired from auditory experience of a tutor song early in life (Tchernichovski et al., 2001). As with human motor skills, song is learned by extensive practice and incremental improvement. Song learning requires a discrete neural ‘song system’ that contains DA-BG-cortical loops similar to mammals (Figure 1B) (Jarvis, 2007; Reiner et al., 2004). Interestingly, the striatopallidal nucleus Area X is the only known BG nucleus of the song system; thus the songbird has not appeared to precisely replicate the dorsal/ventral BG subdivision that may implement AC in mammals (Jarvis, 2007; Reiner et al., 2004). One reason is that song is a learned motor sequence without primary reinforcements, and ventral BG circuits are widely implicated in hedonic processes such as foraging, reward processing, and drug addiction (Apicella et al., 1991; Atallah et al., 2014; Haber and McFarland, 1999; Humphries and Prescott, 2010; Schultz et al., 1992; Tachibana and Hikosaka, 2012; Tindell et al., 2006).

Yet in other respects, parts of the song system appear to embody the AC architecture (Doya and Sejnowski, 1995). The premotor cortical nucleus HVC can generate state-like information in the form of what time-step it is in the song (Hahnloser et al., 2002; Mackevicius and Fee, 2018a). VTA DA neurons resemble the critic’s output; they signal performance prediction error, the difference between how good a syllable just sounded and how good it was predicted to sound based on recent performance (Figure 1C) (Gadagkar et al., 2016). Area X has properties of the actor; it receives state (song time-step) information from HVC as well as dopaminergic prediction error from VTA, and it sends outputs to a premotor thalamocortical circuit capable of driving real-time changes in vocal output (Fee and Goldberg, 2011; Kao et al., 2005; Luo et al., 2001). Further, DA signaling inside Area X modulates corticostriatal plasticity (Ding and Perkel, 2004), is required for learning (Hoffmann et al., 2016), and can reinforce the specific way a syllable is produced (Hisey et al., 2018; Xiao et al., 2018).

The identification of performance prediction error signaling by DA neurons placed renewed focus on inputs to VTA that signal the predicted and actual error of song syllables, i.e. that may complete the role of the critic. One projection to VTA comes from a high-order auditory cortical area, the ventral intermediate arcopallium (AIV) that appears to signal ‘actual’ auditory error, analogous to ‘actual’ reward but with reversed sign (Mandelblat-Cerf et al., 2014). VTA-projecting AIV neurons exhibit low baseline firing rates and are phasically activated by distorted auditory feedback (DAF) during singing. DAF, though not generally aversive (Murdoch et al., 2018), induces a perceived error on distorted renditions such that undistorted renditions are reinforced (Ali et al., 2013; Andalman and Fee, 2009; Hamaguchi et al., 2014; Keller and Hahnloser, 2009; Tumer and Brainard, 2007). During singing, DAF causes activation of VTA projecting AIV neurons and, at a slightly longer latency, pauses in Area X projecting VTA neurons (VTAx), suggesting that AIV can drive pauses in VTA, but it remains unclear whether and how AIV activity affects VTAx firing.

A second input to VTA comes from a ventral striatopallidal region outside the traditional song system (Gale and Perkel, 2010; Gale et al., 2008). We hypothesized that this vBG region may function as a part of the critic. This hypothesis makes several predictions: (1) vBG lesions should disrupt song learning; (2) vBG should receive inputs from brain areas conveying information about state (e.g. precise time-step of the song) as well as information about performance error; (3) vBG neurons should exhibit predicted error signals, for example by producing rate modulations before time-steps of the song previously targeted with DAF; and (4) vBG neurons should send predicted error signals to VTA, which should project back to both vBG and Area X (e.g. pink lines, Figure 1A).

Here we combine lesions, viral tract tracing and electrophysiology to confirm all of these predictions. We show that the vBG-VTA pathway resembles the critic in classic AC networks. Our findings demonstrate that a circuit motif that implements trial and error learning in machines and during reward-seeking in mammals can solve the problem of performance benchmarking during motor sequence learning in songbirds.

## RESULTS

### An auditory cortical projection suppresses VTAx neurons through feedforward inhibition

To identify upstream pathways that may drive dopaminergic error signals, we injected retrograde tracer into the Area X-projecting part of VTA. Consistent with past work, retrogradely labeled neurons were observed in AIV (Gale et al., 2008; Mandelblat-Cerf et al., 2014)(Figure S1). Previous studies showed that VTA-projecting AIV neurons (AIVvta) recorded during singing are activated by errors in auditory feedback (Mandelblat-Cerf et al., 2014), at the same time that VTAx neurons are suppressed (Gadagkar et al., 2016). We wondered if AIV activation could directly induce suppression in VTAx firing. We recorded VTA neurons in anesthetized birds as we electrically stimulated AIV (Figure 1B). AIV stimulation induced phasic suppressions in wide spiking, antidromically identified VTAx neurons and bursts in thin-spiking VTA interneurons that could inhibit VTAx activity (Figure 1, D to E). Thus VTA contains a local inhibitory circuit that can invert excitatory signals from AIV (Figure 1I), consistent with the idea that performance error-induced activations in AIV can drive pauses in VTAx firing during singing. Notably, the AIV projection resembles cortical projections to VTA in mammals that also target local GABAergic interneurons and inhibit dopaminergic firing (Beier et al., 2015; Carr and Sesack, 1999; Moreines et al., 2017; Patton et al., 2013).

### Ventral basal ganglia lesions impair song learning

Following tracer injection into VTAx, retrogradely labeled neurons were also observed in the ventral pallidum and overlying ventromedial striatum (Figure S2), consistent with previous studies (Gale and Perkel, 2010; Gale et al., 2008). Because striatal and pallidal cell types can be spatially intermingled in birds (Person et al., 2008), we term this region ventral basal ganglia (vBG). To test its role in song learning, we performed sham or real excitotoxic vBG lesions in juvenile birds and evaluated their adult songs (STAR Methods). vBG lesions significantly impaired song learning (Song imitation score, paired t test between control and lesioned siblings, p=0.0416; rank-sum test between all controls and all lesioned birds, p=0.014. N=6 lesion, 7 controls. STAR Methods) (Figure 2).

**Figure 2.**
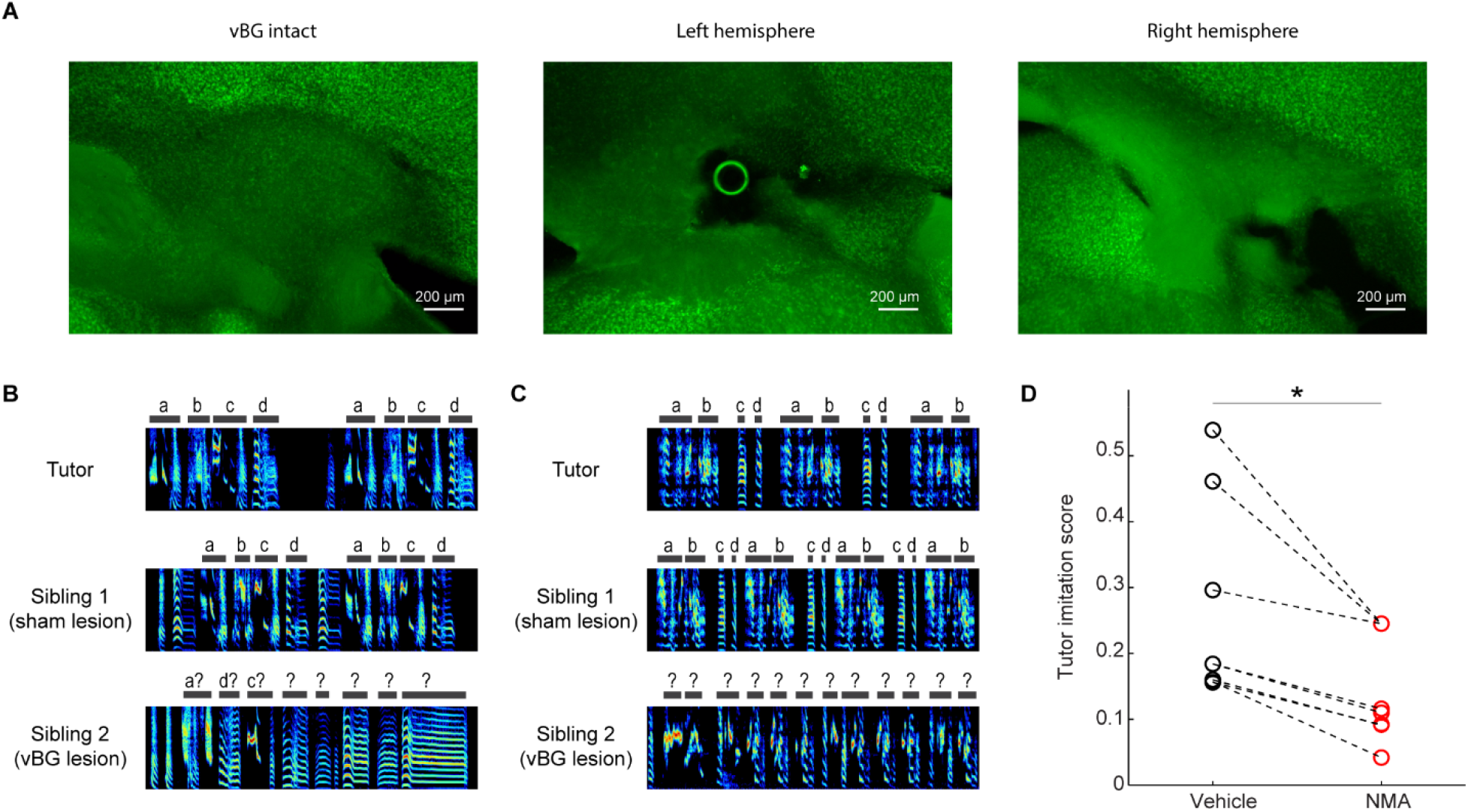
Ventral basal ganglia lesions impair song learning. (A) Lesions were confirmed in neuronal nuclear stained (anti-NeuN) slices as extensive tissue damage and cell death. (B) Tutor song (top), adult song of sham lesioned (middle), and vBG lesioned (bottom) siblings. (C) Same as (B) for another pair. (D) Adult song of vBG lesioned birds had lower similarity to their tutor compared to controls (paired t test, p = 0.0416; rank-sum test between all controls and all lesioned birds, p = 0.014. N=7 sham lesion, N=6 vBG lesion).

### vBG neurons exhibit error signals during singing

Learning deficits following vBG lesion suggest a role in song evaluation. To test how vBG may guide learning, we recorded vBG neurons in singing birds while controlling perceived error with DAF (Figure 1C) (Andalman and Fee, 2009; Tumer and Brainard, 2007)(STAR Methods). Beginning days prior to recordings, a specific ‘target’ song syllable was either distorted with DAF or, on randomly interleaved renditions, left undistorted (distortion rate 48.0±1.4%, mean□±□s.e.m., n=38 birds). Significant auditory error responses were observed in 29/122 vBG cells (Figure 3). We defined responses as error-activated (n=14) or error-suppressed (n=15) (STAR Methods). Some error-suppressed neurons exhibited significant and precisely timed phasic activations immediately following undistorted renditions of the target syllable, analogous to the positive prediction error signal previously observed in VTAx neurons (n=6, Figure 3B) (STAR Methods). Latencies and durations of error responses were commensurate with those observed in downstream VTAx DA neurons (latency: 59.4±4.6 ms, duration: 78.5±7.5ms) (Figure S3). Analysis of movement patterns with microdrive-mounted accelerometers demonstrated that error responses were not attributable to body movement (Figure S4). Error responses were rarely observed following passive playback of song to nonsinging birds (p>0.05 in 13/14 neurons, WRS test), consistent with singing-related performance error (Figure S5, A and B). Yet two neurons appeared purely auditory in nature because they exhibited similar song-locked firing patterns during active singing and passive playback (Figure S5, C and D).

**Figure 3.**
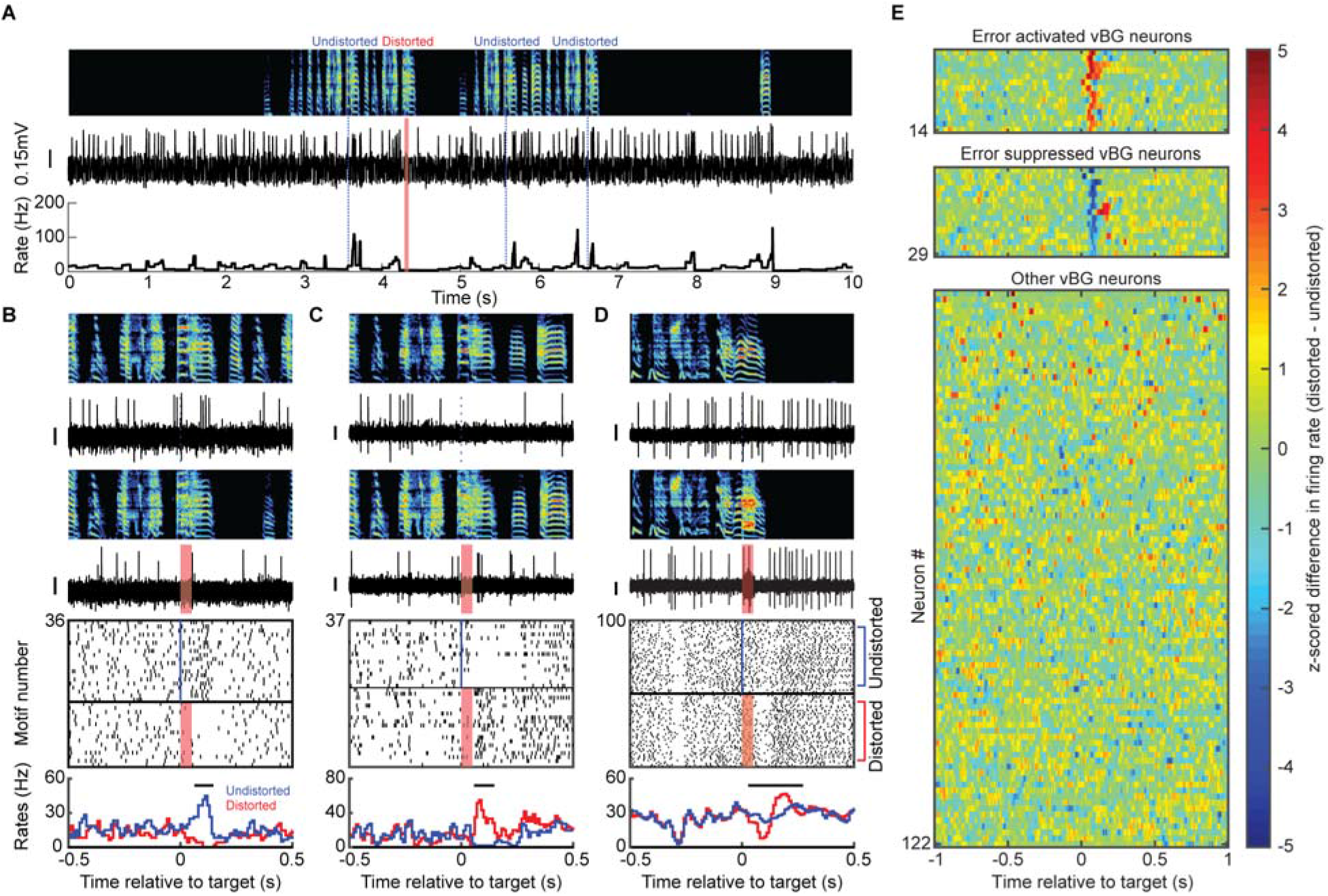
vBG neurons exhibit error responses during singing. (A) Spectrogram (top), discharge (middle) and instantaneous firing rate (bottom) of a vBG neuron recorded during singing. (DAF, red shading; undistorted targets, blue lines). (B) Expanded view of neuron from (A). Top to bottom: spectrograms, spiking activity during undistorted and distorted trials, corresponding spike raster plots and rate histograms (all aligned to target onset). Horizontal bars in histograms indicate significant different response (P < 0.05, WRS test) (STAR Methods). (C and D) Additional examples of error activated and error suppressed neurons, same format as (B). (E) Each row plots the z-scored difference between undistorted and distorted target-aligned rate histograms. Error activated neurons (top, n=14), error suppressed neurons (middle, n=15), and non-error responsive neurons (bottom, n=93) are independently sorted by maximal z-score.

### vBG neurons exhibit temporally precise song-locked activity during singing

To generate precisely timed bursts in positive-prediction error encoding neurons (e.g. Figure 3B), the vBG region requires information about song timing. Many non-error responsive vBG neurons exhibited temporally precise song-locked firing (Figure 4 and Figure S6A, n=8/159, intermotif correlation coefficient>0.3, STAR Methods). One class was distinguished by its ultra-sparse discharge aligned to specific song syllables (Figure 4A and Figure S6B, n=2, sparseness index>0.5, STAR Methods). These neurons’ discharge strongly resembled striatal medium spiny neurons (MSNs) previously recorded in Area X (Goldberg and Fee, 2010; Woolley et al., 2014). Another class was distinguished by stereotyped firing patterns visible as high frequency bursts aligned to specific song time-steps with millisecond precision (Figure 4B and Figure S6C, n=6). Other vBG cell types exhibited prominent phasic activations or suppressions immediately prior to the target-time of the song, consistent with predicted error and predicted quality signals, respectively (Figure 4C and Figure S6, D and E, n=10).

**Figure 4.**
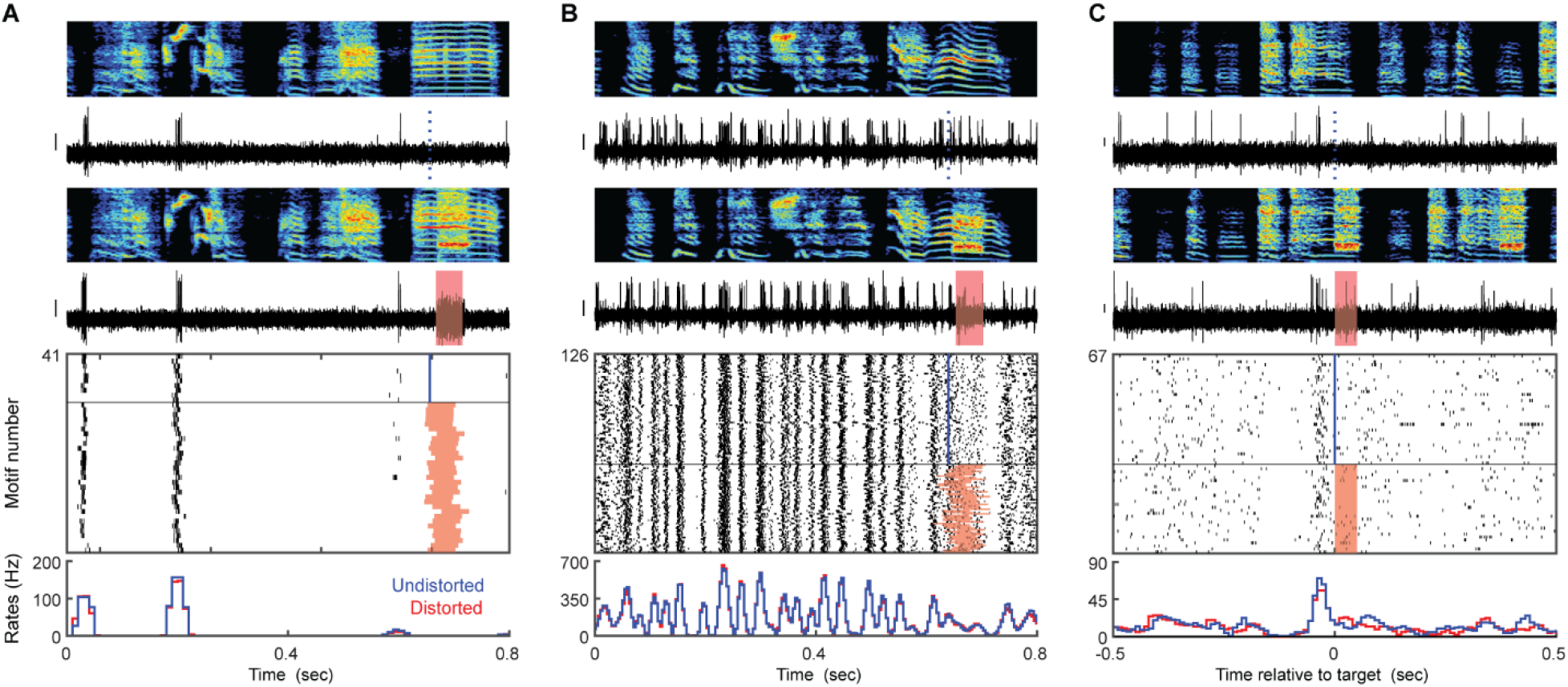
Example vBG neurons that encode precise song timing and pre-target bursts during singing. (A to C) Top to bottom: spectrograms, spiking activity during undistorted and distorted trials, corresponding spike raster plots and rate histograms for a vBG neuron with sparse, temporally precise discharge (A), one with time-locked bursts that tile the song (B), and one with a pretarget burst characterized by pronounced activation immediately prior to the targeted time in the song (C). Y scale bar for spiking activity is 0.15 mV.

Yet other neurons were distinguished by dramatic increase or decrease in firing rate at the transition from non-singing to singing (Figure S6, G to I, n=10, rate difference >85%) or exhibited no detectable rate modulation during singing (Figure S6F, n=60, p>0.01, Kolmogorov-Smirnov test against shuffled distribution, STAR Methods). All of these diverse cell types were spatially intermingled (Figure S7), demonstrating that, unlike other nuclei of the traditional song system, singing and non-singing related neurons are intermixed in the vBG.

### Antidromically identified VTA-projecting vBG neurons exhibit prediction and performance error related activity

If the vBG functions as part of the critic, it should send predicted error-related signals to VTA. We used antidromic and collision testing methods to identify VTA-projecting vBG (vBGvta) neurons (n=7, 100 tested). VTA-projecting neurons with low mean firing rates (<50 Hz) exhibited significant phasic activation following undistorted targets (Figure 5C), consistent with a prediction error signal, or prior to the song target time (Figure 5D) (STAR Methods), consistent with a predicted error signal. vBGvta neurons with high mean firing rates (>50 Hz) exhibited auditory error responses (Figure 5, E and G) and pauses in firing immediately prior to the target time-step of the song (Figure 5, F to H), consistent with a predicted quality signal.

**Figure 5.**
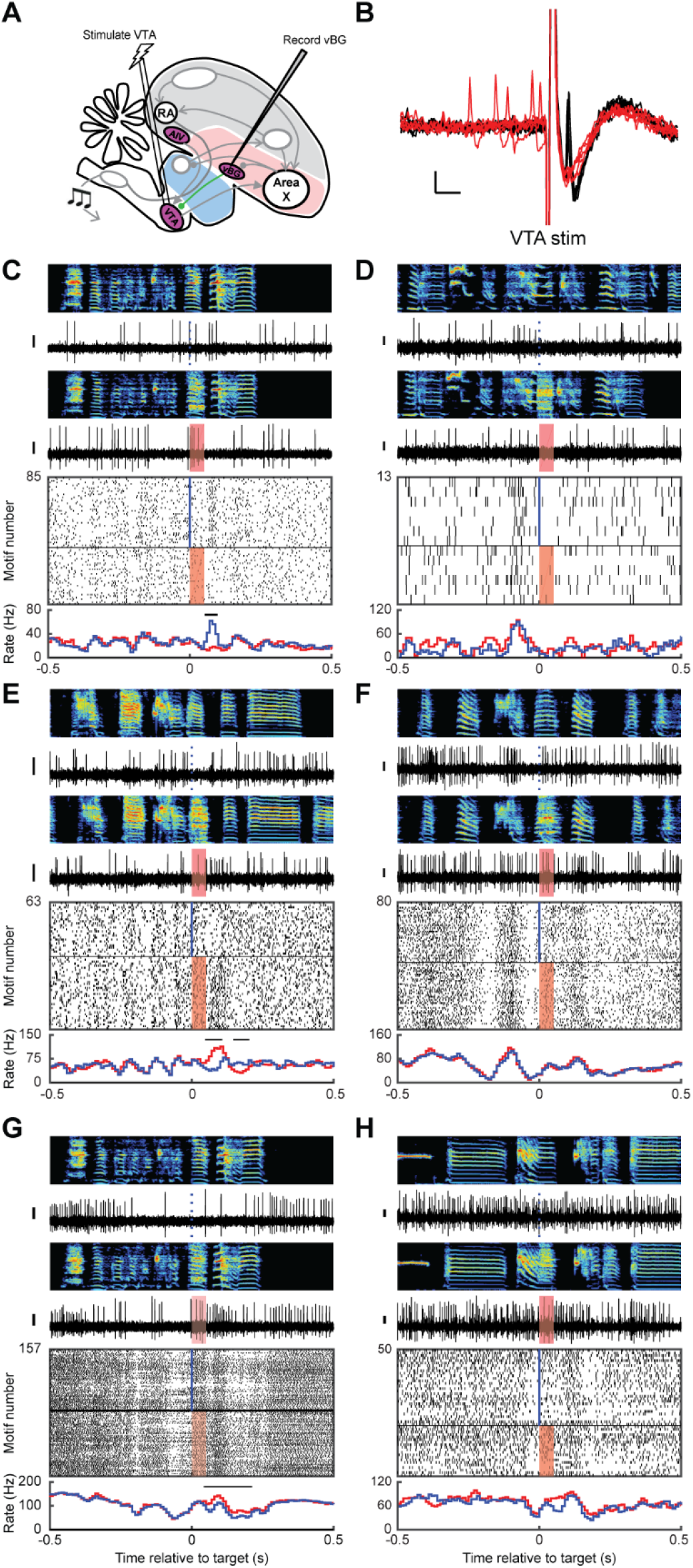
Antidromically identified VTA-projecting vBG neurons exhibit predicted and actual error responses. (A and B) Stimulation and recording electrodes were chronically implanted into VTA and vBG, respectively, for antidromic identification of VTA-projecting vBG neurons (vBGvta). (B) Antidromic (black) and collision (red) testing of the neuron shown in (C). (C to H) Song-locked firing patterns of six confirmed vBGvta neurons, plotted as in Figure 3, reveal activity patterns including activation following undistorted targets (C), pre-target bursts (D), error-induced activation (E), and pre-target pauses (F to H).

### Viral tracing identifies inputs to vBG from vocal motor and auditory regions, and VTAx

What inputs to vBG could account for this stunning diversity of singing, auditory error, and error prediction-related firing? Using retrograde and anterograde viral tracing strategies, we identified inputs to vBG from (1) RA, a vocal motor cortex-like nucleus known to send precise motor command signals to brainstem motor neurons (Leonardo and Fee, 2005; Sober et al., 2008; Yu and Margoliash, 1996); (2) Uva, a motor thalamic nucleus known to send precise song timing information to HVC, a premotor cortical nucleus (Danish et al., 2017; Hamaguchi et al., 2016); (3) DLM, the Area X-recipient thalamic nucleus known to send premotor signals to cortical nucleus LMAN (Goldberg and Fee, 2012); (4) AIV, an auditory cortical area known to send ‘actual’ (just-heard) auditory error signals to VTA (Mandelblat-Cerf et al., 2014); (5) Ovoidalis, the primary auditory thalamus (Lei and Mooney, 2010; Vates et al., 1996); and (6) VTAx neurons, known to be dopaminergic and to send performance prediction error signals to Area X (Gadagkar et al., 2016; Person et al., 2008) (Figures 6 and 7).

**Figure 6.**
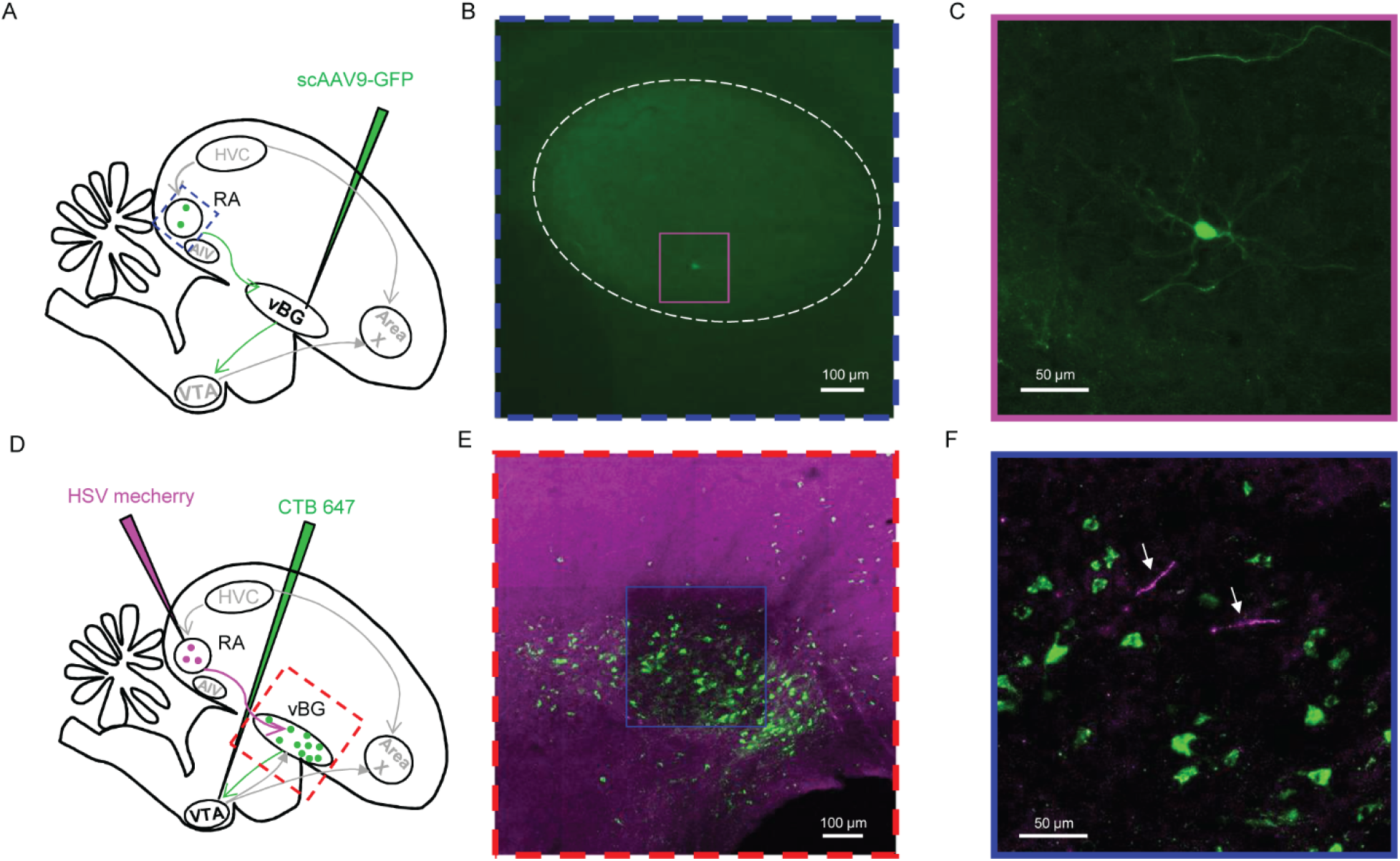
RA projects to vBG. Combined anterograde and retrograde strategies were used to confirm RA projection to vBG. (A) Sparse retrograde virus (scAAV9-GFP) was injected into vBG. (B) Fluorescently labeled RA neuron, centered in pink box (dashed white lines denote RA boundaries). (C) Expanded view of the pink box from (B) revealing GFP-expressing vBG-projecting RA neuron. (D) In anterograde strategy, anterograde virus (HSV-mCherry) was injected into RA and CTB-647 was injected into VTA. (E) Photomicrograph of vBG region, with VTA-projecting neurons clearly visible (green). (F) Expanded view from blue box in (E), revealing proximity of RA axons (purple) to VTA-projecting neurons in vBG.

**Figure 7.**
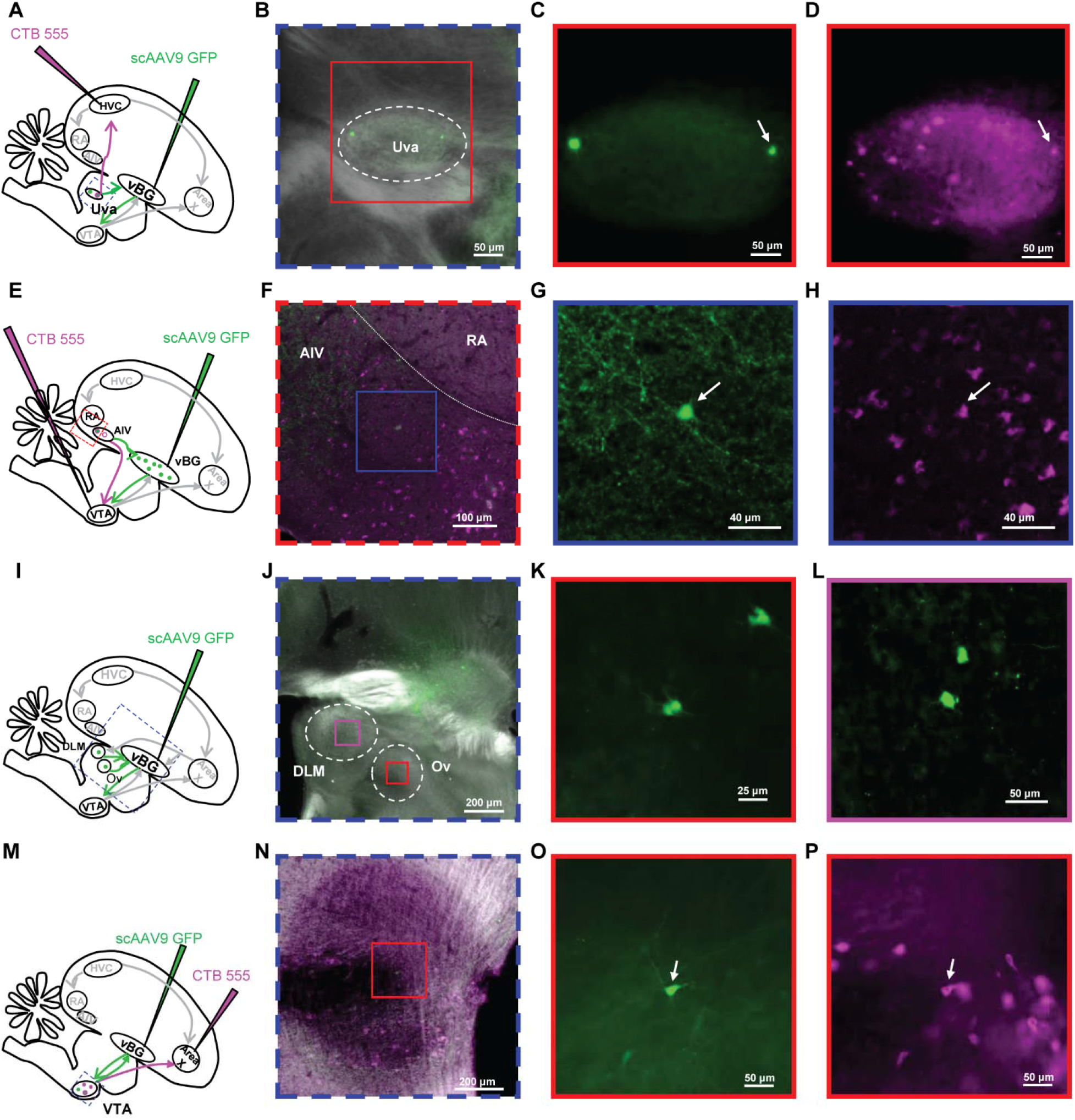
Uva, AIV, DLM, Ov and VTAx project to vBG. To label vBG-projecting neurons, sparse retrograde scAAV9 virus was injected into vBG (A,E,I,M). (A to D) HVC-projecting nucleus Uvaeformis (Uva) thalamic neurons project to vBG. CTB-555 was injected into HVC (A). Retrogradely labeled vBG-projecting Uva neuron (arrow, C) was colabeled with HVC-projecting Uva neuron (arrow, D). (E to H) VTA-projecting AIV neurons also project to vBG. CTB-555 was injected into VTA (E). Retrogradely labeled vBG-projecting AIV neuron (arrow, G) is colabeled with VTA-projecting AIV neuron (arrow, H). (I to L) Area X recipient thalamus DLM and primary auditory thalamus Ovoidalis (Ov) project to vBG. Retrogradely labeled DLM neurons (K) and Ov (L). (M to P) Area X-projecting VTA dopamine neurons (VTAx) send collaterals to vBG. CTB-555 was injected into Area X (M). Retrogradely labeled vBG-projecting VTA neuron (arrow, O) was colabeled with X-projecting VTA neuron (arrow, P).

## DISCUSSION

By combining lesions, viral tract tracing, and electrophysiology we discovered that vBG is required for song learning, receives information about song time-step and auditory error, generates predicted and prediction error signals, and sends them to VTA. These findings expand the avian song system and, more generally, identify a critic component of an AC-like network. AC networks, commonly deployed in machine learning and implicated in reward processing in mammals, can thus implement performance evaluation during a purely motor sequence learning task like birdsong.

Our results suggest a general principle for how to compute prediction error. Specifically, the reciprocal vBG-VTAx loop we identified strongly resembles ‘spiraling’ pathways that link ventral to dorsal BG through VTA in mammals, a cornerstone of actor-critic models of prediction error computation and reward-based decision making (pink lines, Figure 1A and 8) (Daw et al., 2006; Haber et al., 2000; Joel et al., 2002; Takahashi et al., 2008). In these models, a ventral critic implements DA-modulated corticostriatal plasticity to learn a state-dependent value function, for example a time-, place-, or cue-dependent reward prediction. This ventral critic projects to VTA and provides DA neurons with the temporally precise ‘prediction’ component of reward prediction error (Takahashi et al., 2016). VTA dopamine neurons project back to the ventral critic to update predicted state-value as well as to a dorsal actor which implements DA-modulated corticostriatal plasticity to learn a reward-maximizing policy. Such AC architectures were first implemented in machines and effectively guide skill and game learning (Gullapalli et al., 1994; Peng et al., 2016; Silver et al., 2016; Sutton and Barto, 1998; Tesauro).

Invoking the AC architecture provides explanatory power for understanding the vBG connectivity and neural signals we observed (Figure 8B). Specifically, thalamic (Uva) inputs could provide state representations in the form of ‘time-step’ in song (Danish et al., 2017) that could explain the observed vBG timing responses (Figure 4). AIV inputs provide information about ‘actual’ (just-heard) auditory error and could explain the error responses (Figure 3) (Mandelblat-Cerf et al., 2014). VTAx inputs could enable DA-modulated thalamostriatal plasticity of Uva inputs to compute predicted state-value in the form of predicted syllable quality (Figure 8B). This explains the error prediction signals such as the pre-target bursts and pauses that can be routed from vBG to VTA (Figure 5, D and F to H). Thus, the vBG contains information necessary to signal the difference between predicted and actual error, manifest in the observed prediction error responses (Figure 3, A and B) that can be sent to VTA (Figure 5C). As in mammals, dopaminergic prediction error signals project to vBG where they could update predicted state-value, as well as to a dorsal BG module (Area X) where they could update the action policy, i.e. how to sing a given syllable (Fee and Goldberg, 2011). Consistent with this view, manipulation of DA activity in Area X can reinforce the way a target syllable is produced (Hisey et al., 2018; Xiao et al., 2018), much like manipulation of striatal DA in mammals can reinforce place preference or action selection (Corbett and Wise, 1980; Tsai et al., 2009; Wise and Schwartz, 1981).

**Figure 8.**
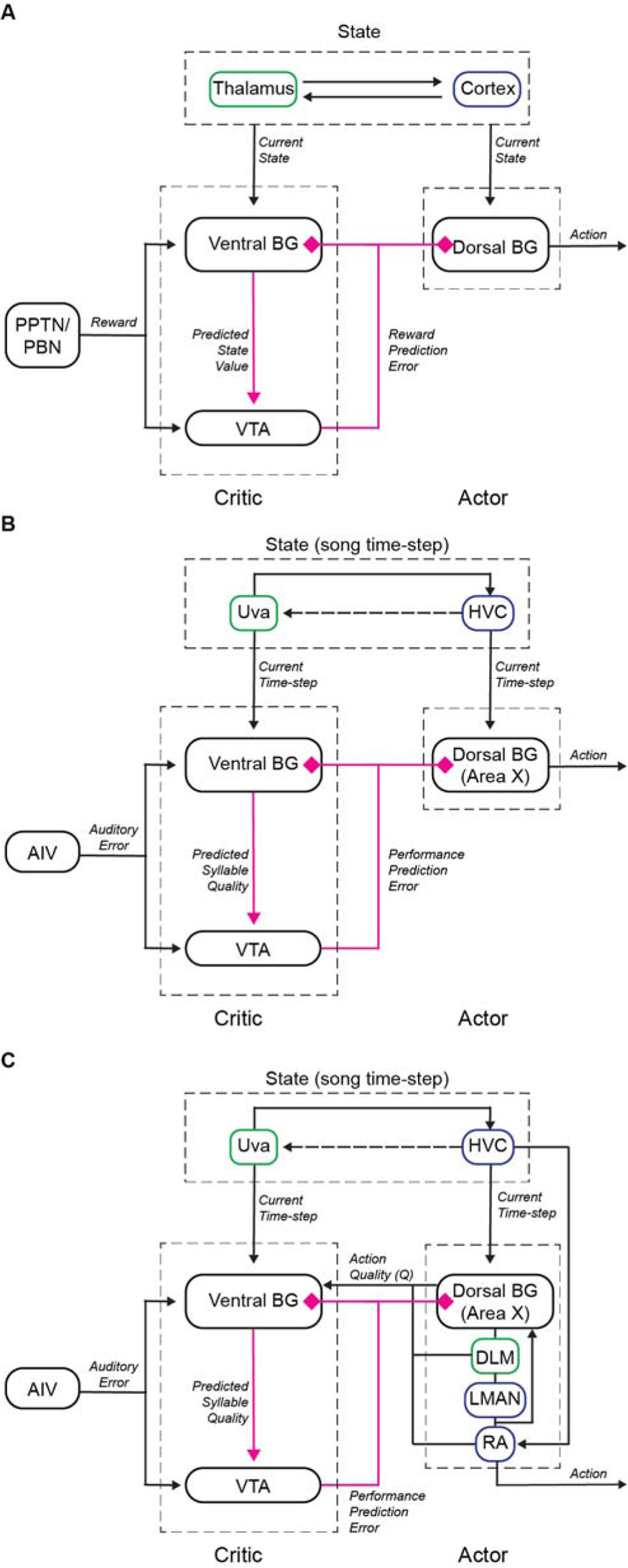
Possible actor-critic circuit motifs for computing predicted performance quality. (A) Actor-critic circuit motif in mammalian BG inspired by refs (Joel et al., 2002; Sutton and Barto, 1998; Takahashi et al., 2008). The critic computes an RPE signal based on the difference between a state-dependent reward prediction and that actual reward received. (B-C) Anatomy and signaling in songbird vBG reveals a similar motif. (B) In this model, vBG learns the predicted syllable quality, and Area X learns to produce the performance maximizing action. The critic implements DA-modulated plasticity of Uva (‘state’) inputs to compute error-weighted timing or, equivalently, song-timestep dependent predicted error. The critic also computes the performance prediction error by comparing the predicted error to actual error (signaled by AIV). (C) An advantage-actor-critic inspired network that additionally makes use of the Area X, DLM and RA inputs to vBG. These inputs enable the critic to compute the advantage function, the predicted quality of a specific vocal variation compared to the average quality at a specific time-step. As in B, this prediction is compared with the actual error, signaled by AIV, to compute the dopaminergic performance prediction error.

Considering how neural circuits implement RL requires careful specification of what constitutes ‘state’ and ‘action’ (Barto, 1995). In standard RL architectures, state representations must access all possible action representations because before learning it is not yet known which state/action pair will produce reward (Joel et al., 2002; Suri and Schultz, 1998). Therefore, a neuron that represents a state should project to all motor channels. In contrast, a neuron that projects to specific motor channels and not others cannot represent state and is well suited to play the actor. In motor systems, many structures have conserved topographical projection patterns that eventually innervate specific muscles; neurons in these structures are candidate ‘actors’. Yet other structures have non-topographic outputs that can contact many motor channels; neurons in these structures provide candidate ‘state’ representations. Note that the critic’s output must also be non-topographic, as it must be able to modify the synaptic strength of all possible state/action pairs.

Based on these considerations, Area X is well suited to constitute part of the actor because it projects topographically through DLM, LMAN, RA and brainstem regions that ultimately innervate distinct muscles of the vocal organ (Fee and Goldberg, 2011; Luo et al., 2001; Vicario and Nottebohm, 1988; Wild, 1993). Put simply, each of these structures can be imagined as a piano keyboard, in which the position of a neuron relates to the muscles it will innervate. Consistent with this idea, neuronal activities in this pathway drive specific actions over others, e.g. how to sing a given syllable (Hisey et al., 2018; Kao et al., 2005; Sober et al., 2008; Xiao et al., 2018). Meanwhile, Uva and HVC are well suited to provide state-like representations. Uva’s access to vocal motor output goes through HVC, which projects non-topographically to RA and Area X (Luo et al., 2001), i.e. the axon of a single HVC neuron can traverse the whole piano. Time-step information in the Uva-HVC pathway provides a sensible state representation for a learned motor sequence. Just as each location in a foraging task has its own history of reward, each time-step of a motor sequence has its own history of error. Similarly, just as a foraging policy specifies which way to go at a given place in a landscape, song policy specifies what note to sing at a given time-step in the sequence.

Curiously, we found that parts of the proposed actor pathway (Area X, DLM and RA) project to vBG (Figures 6–7)(see also (Gale et al., 2008)), revealing a non-standard projection from actor back to critic. This may allow the critic to calculate a refined prediction based not only on the state, but also on the action taken. Notably, a growing family of AC algorithms could make use of such projections (Mnih et al., 2016; Schulman et al., 2015). In the classic AC algorithm described above (Figures 1, 8A,B), the critic’s RPE signal derives from the difference between the reward received, r, and the reward predicted based on the state, V(s_t_). In equation 1, V is the value of a state, defined as the expected total reward in future. Predictions are updated by the difference between predicted and actual reward using temporal difference (*δ*) (equation 2).

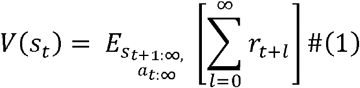

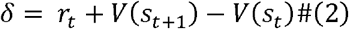

In ‘advantage-actor-critic’ (A2C) networks, the critic is subtly different. It evaluates actions by additionally computing the quality, Q(s_t_,a_t_), of a specific state/action pair (equation 3). The reinforcement signal derives from the difference between Q and the average value of the state, a quantity called the ‘advantage’ (equation 4). Advantage values are updated by a prediction error based on the temporal difference (equation 5) (Dayan and Balleine, 2002; Mnih et al., 2016; Schulman et al., 2015).

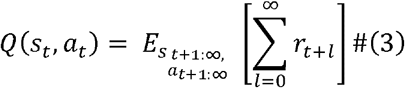

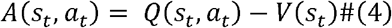

'

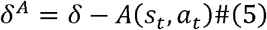

Here, only state/action pairs that produce outcomes that exceed the average state value are reinforced; this scales policy evaluations and reduces their variance (Mnih et al., 2016; Schulman et al., 2015). Notably, in contrast to standard AC networks, the critic of the A2C network requires Q to be approximated, which in turn requires information about action as well as state and outcome (Figure 8C).

Thus Area X, DLM and RA inputs to vBG raise the possibility that the songbird could compare actual song quality not just to the quality predicted at that time-step, but additionally to the predicted quality of the action taken at that time-step, analogous to what an A2C network does. A simple experiment to test this possibility would be to record VTAx DA neurons while manipulating the advantage, A(s_t_,a_t_). This could be done by implementing conditional distorted auditory feedback in which only low-pitch variants of a specific target syllable are distorted (Andalman and Fee, 2009; Tumer and Brainard, 2007). On rare catch trials, low pitch variants would instead be left undistorted. If DA neurons signal the difference between the actual outcome to the predicted outcome given the state (the target time of the song), then the magnitude of DA bursts would be the same for all undistorted target renditions, regardless of which syllable variant was produced. But if DA neurons signal the advantage, then the DA burst following the undistorted rendition of the low pitch variants will be larger than following the high pitch renditions, because low pitch renditions have been associated with histories of more error. Future recordings of VTAx neurons could therefore constrain which variant of AC-like algorithms is realized in the songbird.

We further speculate that Area X could compute Q both in its role as actor and as an input to the critic in an A2C network. In birdsong, Q(s,a) is the predicted action quality of a given vocal variation at a specific state, or song timestep. DA-modulated plasticity of HVC inputs to Area X (Ding and Perkel, 2004) can in principle weigh the value of all state/action pairs (i.e. compute Q) because of the unique anatomy of Area X: its inputs from HVC and VTA are non-topographic and yet its projections to DLM are segregated into topographically organized output ‘channels’ (Luo et al., 2001). This idea also makes specific experimental predictions. If performance at a given time-step is poor no matter what action is taken, then the predicted quality of all state/action pairs will be low. This occurs naturally during vocal babbling, before the onset of learning, when the predicted quality of all syllables is likely to be low. At this learning stage, all Area X output neurons exhibit phasic activations that are concentrated at syllable onsets, exactly where state representations (HVC activity) are also clustered (Okubo et al., 2015; Pidoux et al., 2014). We hypothesize that this is Area X’s way of saying it has not yet learned any policy to promote. These pre-syllable activations rapidly go away over days of singing experience, as birds have an opportunity to learn which state/action pairs produce good outcomes (Goldberg et al., 2010; Goldberg and Fee, 2010). This hypothesis also predicts that all Area X output neurons should exhibit rate increases prior to a song time-step targeted with DAF with 100% probability.

Many open questions remain. Foremost, it remains unclear how AIV computes its auditory error signal. Extracting this error signal may require comparison with tutor song, which appears to be stored in other auditory cortical areas (Mackevicius and Fee, 2018b). It may also require comparison of actual acoustic signals against predicted acoustic signals, which could rely on a forward model of song production (e.g. state-dependent sound prediction) implemented by HVC projections to the auditory system (Ganguli and Hahnloser, 2011; Roberts et al., 2017). It also remains unclear how Area X outputs are transformed into action by downstream circuits. Finally, whereas our findings emphasize BG circuits, song learning also requires plasticity in cortical nuclei of the song system (Ali et al., 2013; Andalman and Fee, 2009). Adult song remains intact following lesions of the Area X→LMAN pathway, suggesting that the song motor program is stored in the HVC-RA projection (Scharff and Nottebohm, 1991). One idea is that action quality encoding by Area X→LMAN results premotor bias that is consolidated into HVC-RA synapses by heterosynaptic Hebbian mechanisms (Andalman and Fee, 2009; Garst-Orozco et al., 2014). Notably, cholinergic signaling in RA regulates synaptic plasticity and is required for vocal learning (Puzerey et al., 2018; Salgado-Commissariat et al., 2004), and cholinergic projections to RA arise from the same vBG region we have studied (Gale et al., 2008). It will be interesting to test if the vBG→RA projection, which is analogous to ascending cholinergic projections in mammals, also encode error or prediction signals useful for learning.

## ACKNOWLEDGEMENTS

We thank Todd Roberts for providing virus, Joe Fetcho, Teja Bollu, and Josh Dudman for comments, and Vikram Gadagkar, Aaron Andalman, and Dmitriy Aronov for comments and analysis code. Funding to JHG was provided by the NIH (R01NS094667), the Pew Charitable Trust, and the Klingenstein Neuroscience Foundations. Funding to PAP was provided by the NIH (F32NS098634). Imaging data was acquired in the Cornell BRC-Imaging Facility using the shared, NIH-funded (S10RR025502) Zeiss LSM 710 Confocal.

## AUTHOR CONTRIBUTIONS

RC, PAP, ACR, TER, KM, AP, AF, and JHG acquired data. RC, PAP, AP, KM, and JHG analyzed data. RC and JHG wrote the manuscript.

## STARE Methods

### CONTACT FOR REAGENTS AND RESOURCE SHARING

Further information and requests for resources and reagents should be directed to and will be fulfilled by the Lead Contact, Jesse H. Goldberg

### EXPERIMENTAL MODEL AND SUBJECT DETAILS

#### Subjects

Subjects were 61 male zebra finches. Animal care and experiments were carried out in accordance with NIH guidelines and were approved by the Cornell Institutional Animal Care and Use Committee.

### METHOD DETAILS

#### Surgery and histology

All surgeries were performed with isoflurane anesthetization. For functional mapping experiments (4 birds, Figure 1), bipolar stimulation electrodes were implanted into AIV and Area X(Gadagkar et al., 2016; Mandelblat-Cerf et al., 2014). Briefly, AIV coordinate was determined by its anterior and ventral position to RA, and Area X coordinate was +5.6A, +1.5L relative to lambda and 2.65 ventral relative to pial surface, at a head angle of 20 degrees. Recordings were made in VTA using a carbon fiber electrode (1 MOhm, Kation Scientific). VTA was identified by anatomical landmarks. Specifically, the boundaries of DLM and Ovoidalis were determined by spontaneous firing and auditory responses. Recordings were then made at the same AP position, +0.6L relative to lambda and 6.5 ventral relative to pial surface, at a head angle of 55 degrees. Area X projecting neurons were further confirmed by antidromic response and collision testing. Location of the stimulating electrodes was verified histologically.

For vBG lesion (13 birds, 39-52 dph), a bipolar stimulation electrode was implanted into Area X and the center of vBG was electrophysiologically mapped by finding units suppressed by Area X stimulation. 115nl of 2% N-methyl-DLaspartic acid (NMA; Sigma, St Louis, MO) was injected into vBG bilaterally. Lesion was verified histologically by anti-NeuN staining.

For awake-behaving electrophysiology (38 birds, 87-355 dph), custom made microdrives carrying an accelerometer (Analog Devices AD22301), linear actuator (Faulhaber 0206 series micromotor) and homemade electrode arrays (5 electrodes, 3-5 MOhms, microprobes.com) were implanted into vBG by coordinates (4.4-5.4A, 1.1-1.5L, 3.5V, head angle 20 degrees). In 19/38 birds, a bipolar stimulation electrode was implanted into VTA using anatomical landmarks as described above. After each experiment, small electrolytic lesions (30 μA for 60 s) were made with one of the recording electrodes. Brains were then fixed, cut into 100 μm thick sagittal sections for histological confirmation of stimulation electrode tracks and reference lesions.

For vBG tracing experiments (6 birds), 40nl of self-complementary adeno-associated virus (scAAV9) with CBh promoter carrying GFP was injected into vBG in two coordinates (4.6/4.9A, 1.3L, 4V). Upstream neurons retrogradely infected and expressing GFP could be observed in RA, AIV, Uva, Ov, DLM, and VTA. To determine if vBG share common inputs with HVC, in addition to scAAV9 in vBG, fluorescently labeled cholera toxin subunit B (CTB, Molecular Probes) was injected into HVC. To determine if vBG share common inputs with Area X, CTB was injected into Area X, and brain sections were immuno-stained with antibodies to tyrosine hydroxylase (Millipore AB152, 1:1000).

#### Functional mapping between AIV and VTA

Neurons were classified as Area X-projecting (VTAx) based on antidromic stimulation and collision testing (200 μs pulses, 100-300 μA). A burst of AIV stimulation consisting three 200 μs pulses with 10ms inter-pulse-interval was delivered every 1.5-2 s, with 300 μA current amplitude. VTA neurons not responsive to Area X stimulation were also tested for AIV stimulation. All VTAx neurons and those putative interneurons activated by AIV stimulation were further analyzed. To determine if VTA neurons respond to AIV stimulations, spiking activity within ±1 second relative to stim burst onset was binned in a moving window of 30 ms with a step size of 5 ms. Each bin within after stim was tested against all the bins in the previous 1 second (the prior) using a z-test. Windows where at least 4 consecutive bins with p<0.05 were considered significant. The response onset and offset were required to bracket lowest (for phasic decrease) or highest (for phasic increase) firing rate after stim onset. For the simultaneously recorded putative VTA interneuron (PIN) and VTAx neuron (Figure 1d-g), we constructed rate histogram of VTAx neuron spiking events aligned to spiking events of PIN with preceding ISI > 10ms. To assess the significance of VTAx rate suppressions following PIN spiking, 1000 surrogate rate histograms were generated by randomly time-shifting each trial of PIN spiking aligned data over the duration of the histogram (2 seconds). Response was considered significant when VTAx firing rate dropped below 5th percentile or exceeded 95th percentile of the surrogate data.

#### Song imitation score

Song learning in vBG lesioned and control birds was assessed by song similarity between pupil (at 90 dph) and their tutors. We computed imitation scores using an automated procedure based on Sound Analysis Pro (SAP) algorithm (Mandelblat-Cerf et al., 2014; Tchernichovski et al., 2000). Briefly, the tutor motif was segmented into syllables by hand. Syllables in the pupil song were determined by finding the section of pupil song with highest SAP similarity to each tutor syllable. Additionally, a sequencing score was computed as the similarity of the next syllable in tutor song and the next section in the pupil song. Imitation score was the product of song similarity and sequence similarity(Mandelblat-Cerf et al., 2014).

#### Syllable-targeted distorted auditory feedback

Postoperative birds with microdrive implant were placed in a sound isolation chamber and given at least a day to habituate to distorted auditory feedback (DAF) as described previously(Gadagkar et al., 2016). Briefly, ongoing singing was analyzed by Labview software to target specific syllables, and two speakers inside the chamber played a 50ms DAF sound on top of bird’s singing on 50% of randomly selected target renditions. DAF was either a broadband sound band passed at 1.5-8 kHz, the same spectral range of zebra finch song, or a segment of one of the bird’s own non-target syllables displaced in time.

#### Passive playback of the bird’s own song

For passive playback of the bird’s own song, we played back randomly interleaved renditions of the undistorted and distorted motifs of the bird’s own song during awake, nonsinging periods. The loudness of playback was adjusted to match the average peak loudness of zebra finch song(Gadagkar et al., 2016; Mandelblat-Cerf et al., 2014).

#### Analysis of neural activity

Neural signals were band-passed filtered (0.25-15 kHz) in homemade analog circuits and acquired at 40 kHz using custom Matlab software. Single units were identified as VTA-projecting by antidromic identification and antidromic collision testing (Figure 4A,b). Spike sorting was performed offline using custom Matlab software. Instantaneous firing rates (IFR) were defined at each time point as the inverse of the enclosing interspike interval (ISI). Firing rate histograms were constructed with 10 ms bins and smoothed with a 3-bin moving average. All data was acquired during undirected song, except for the neuron in Figure 5H, which was recorded during female-directed song.

#### Performance error response

To identify performance-error related neurons, we assessed the difference in firing rate between distorted and undistorted singing renditions as previously described(Keller and Hahnloser, 2009; Mandelblat-Cerf et al., 2014). Neurons with less than 10 trials of either distorted or undistorted renditions of the target syllable were excluded from this analysis. Briefly, we performed a WRS test (p=0.05) on the number of spikes in distorted vs. undistorted renditions in 30 ms windows. Windows were shifted in 5 ms steps and considered significant when at least 4 consecutive windows had p<0.05. Error-related neurons were classified as error-activated if the firing rate is higher in distorted trials in window of significance, and error suppressed if the firing rate is higher in undistorted trials. For error suppressed neurons, we also performed z-test on the firing rate following undistorted trials with 500ms before target as baseline period. Neurons with significant rate increases following target were identified as prediction error neurons.

To test if error responses were attributable to purely auditory responses to a different sound, we performed the same analysis for distorted and undistorted renditions during passive playback of bird’s own song (BOS) playback in 12/24 error neurons. Only one neuron exhibited an error response during passive playback. This neuron also exhibited similar song-locked firing during both singing and listening (Figure S5C). One other error responsive neuron also appeared auditory, although the part of playback that contained target syllable was masked by calls (Figure S5D).

We compared the latency and duration of error response to those of VTAx neurons from a previous dataset(Gadagkar et al., 2016). Latency and duration was defined by the onset and onset-offset interval of significant windows in WRS test as described above. Two auditory neurons described above were not included in this analysis.

#### Song timing related activity

Sparseness index was used to identify putative song-related MSNs. This method has been described elsewhere and separates MSNs from other striatal cell types in the dorsal basal ganglia nucleus Area X(Goldberg and Fee, 2010). For each neuron, we calculated rate histograms aligned to syllable onset for all syllables. Then we normalized these histograms over all syllables to generate a probability density function *p_i_* over N bins. An entropy-based sparseness index was computed as follows:

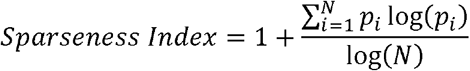

Intermotif pairwise correlation coefficient (CC) was used to identify neurons that had highly time-locked firing to song motifs (timing neurons), as previously described(Goldberg and Fee, 2010). Motif aligned IFR was time warped to the median duration of all motifs, mean-subtracted, and smoothed with a Gaussian kernel of 20ms SD, resulting in *r_i_* for each motif. The Intermotif CC was defined as the mean value of all pairwise CC between *r_i_* as follows:

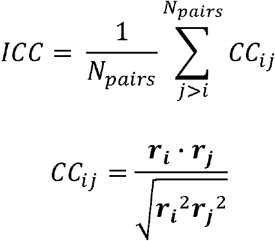

To assess significance of CC distributions, we computed new pairwise CC values for each neuron by adding random, circular time shifts to each spiketrain. CC was considered significant when the real distribution was significantly different from that of randomly shuffled data (P<0.01, Kolmogorov-Smirnov test).

#### Quantification of movement

An accelerometer (Analog Devices AD22301) was mounted on microdrives to quantify gross body movements as described previously(Gadagkar et al., 2016). Briefly, movement onsets and offsets were determined by threshold crossings of the band-passed, rectified accelerometer signal. To test if error responses could be explained by a difference in movement rate following DAF, for each bird we calculated onset times of movements relative to song target time. Then we performed a WRS test (p=0.05) on the number of movement onsets in distorted vs. undistorted renditions in 30 ms windows. Windows were shifted in 5 ms steps and considered significant when at least 4 consecutive windows had p<0.05.

#### Imaging

Imaging data was acquired with a Leica DM4000 B microscope and a Zeiss LSM 710 Confocal microscope. Image processing was done with ImageJ.

### DATA AND SOFTWARE AVAILABILITY

The data that support the findings of this study are available from the corresponding author upon reasonable request.

Key Resources Table

Key Resources Table
| REAGENT or RESOURCE | SOURCE | IDENTIFIER |
| --- | --- | --- |
| Antibodies |  |  |
| Anti-Tyrosine Hydroxylase antibody | Millipore | Cat#AB152 |
| Anti-NeuN antibody, Alexa Fluor 488 conjugated | Millipore | Cat#MAB377X clone A60 |
| Bacterial and Virus Strains |  |  |
| Self-complementary AAV9-CBh-GFP | Gene Therapy Center Vector Core, University of North Carolina at Chapel Hill | N/A |
| Short Term HSV-mCherry | Gene Delivery Technology Core, Massachusetts General Hospital | Cat#RN3 |
| Chemicals, Peptides, and Recombinant Proteins |  |  |
| N-Methyl-DL-aspartic acid | Sigma | Cat#M2137 |
| Experimental Models: Organisms/Strains |  |  |
| Zebra Finch (*Taeniopygia guttata*) | Magnolia Bird Farm, Anaheim CA | N/A |
| Software and Algorithms |  |  |
| MATLAB | MathWorks | https://www.mathworks.com/products/matlab.html |
| ImageJ | NIH | https://imagej.nih.gov/ij/ |
| Sound Analysis Pro | Tchernichovski, O., Nottebohm, F., Ho, C.E., Bijan, P., Mitra, P.P. (2000). A procedure for an automated measurement of song similarity. *Animal Behaviour* 59, 1167-1176 | N/A |
| Similarity Index | Mandelblat-Cerf Y, Fee MS (2014) An Automated Procedure for Evaluating Song Imitation. PLoS ONE 9(5): e96484 | N/A |

## Supplementary Figures

**Figure S1.**
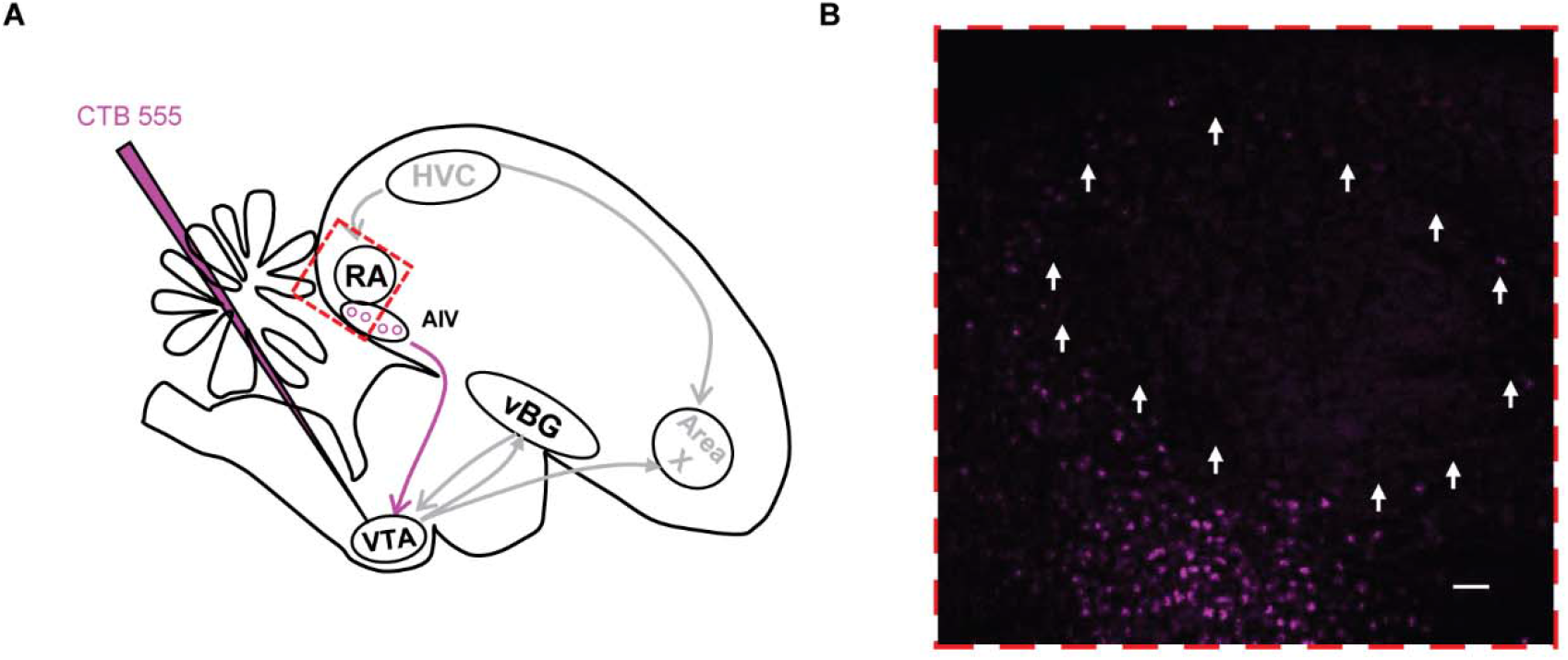
AIV projects to VTA. (A) Schematic of tracing experiment. Injection of CTB-555 into VTA retrogradely labeled cell bodies in AIV, as previously reported(Gale et al., 2008; Mandelblat-Cerf et al., 2014). (B) Expanded view of red square from (A). VTA-projecting neurons are visible in AIV, which surrounds boundaries of RA, denoted by white arrows.

**Figure S2.**
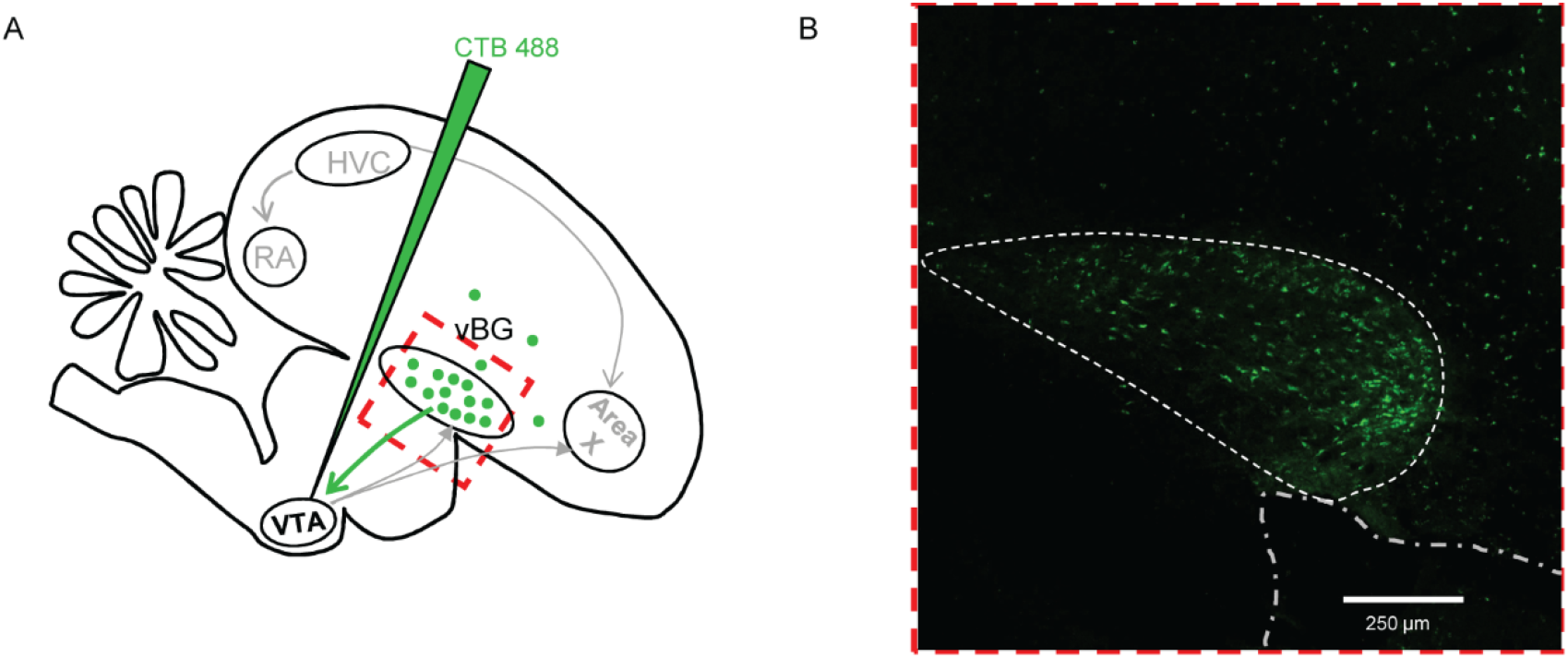
Ventral basal ganglia projects to VTA. (A) Schematic of tracing experiment. Injection of CTB-488 into VTA retrogradely labeled cell bodies in ventral basal ganglia, as previously reported(Gale et al., 2008). (B) Expanded view of red square from (A). VTA-projecting neurons are visible in VP, bounded by dashed white lines, and overlying striatum.

**Figure S3.**
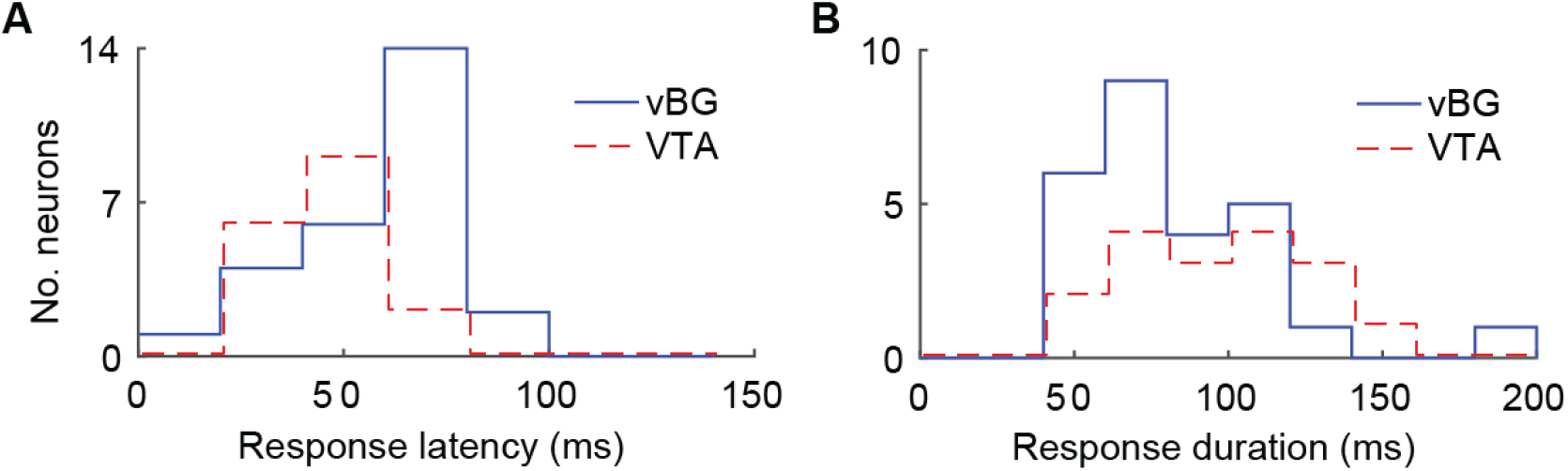
Error response latencies and durations in vBG and VTA neurons. (A) Histograms of response latency for all error responsive neurons. Blue: vBG neurons (n=29, range: 15 ms to 90 ms) Red: VTAerror neurons (n=17, range: 21 ms to 67 ms). (B) Histogram of response duration for all error responsive neurons. Blue: vBG neurons (range: 45 ms to 210 ms) Red: VTA neurons (range: 58 ms to 148 ms). VTA data taken from a previous study (Gadagkar et al., 2016).

**Figure S4.**
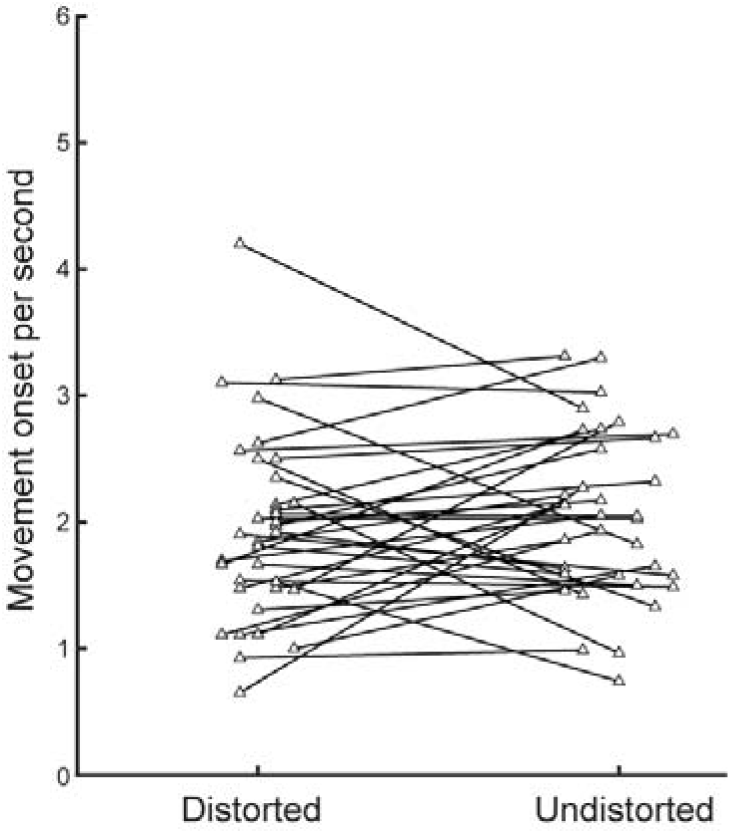
Animal movement was not affected by DAF during singing. Analysis of DAF-related movement responses across all birds. Each line represents data from one bird with average rate of movement onsets in 150 ms following distorted and undistorted syllables. There was no difference in movement between distorted and undistorted motifs (p > 0.05 in 35/35 birds, WRS test.)

**Figure S5.**
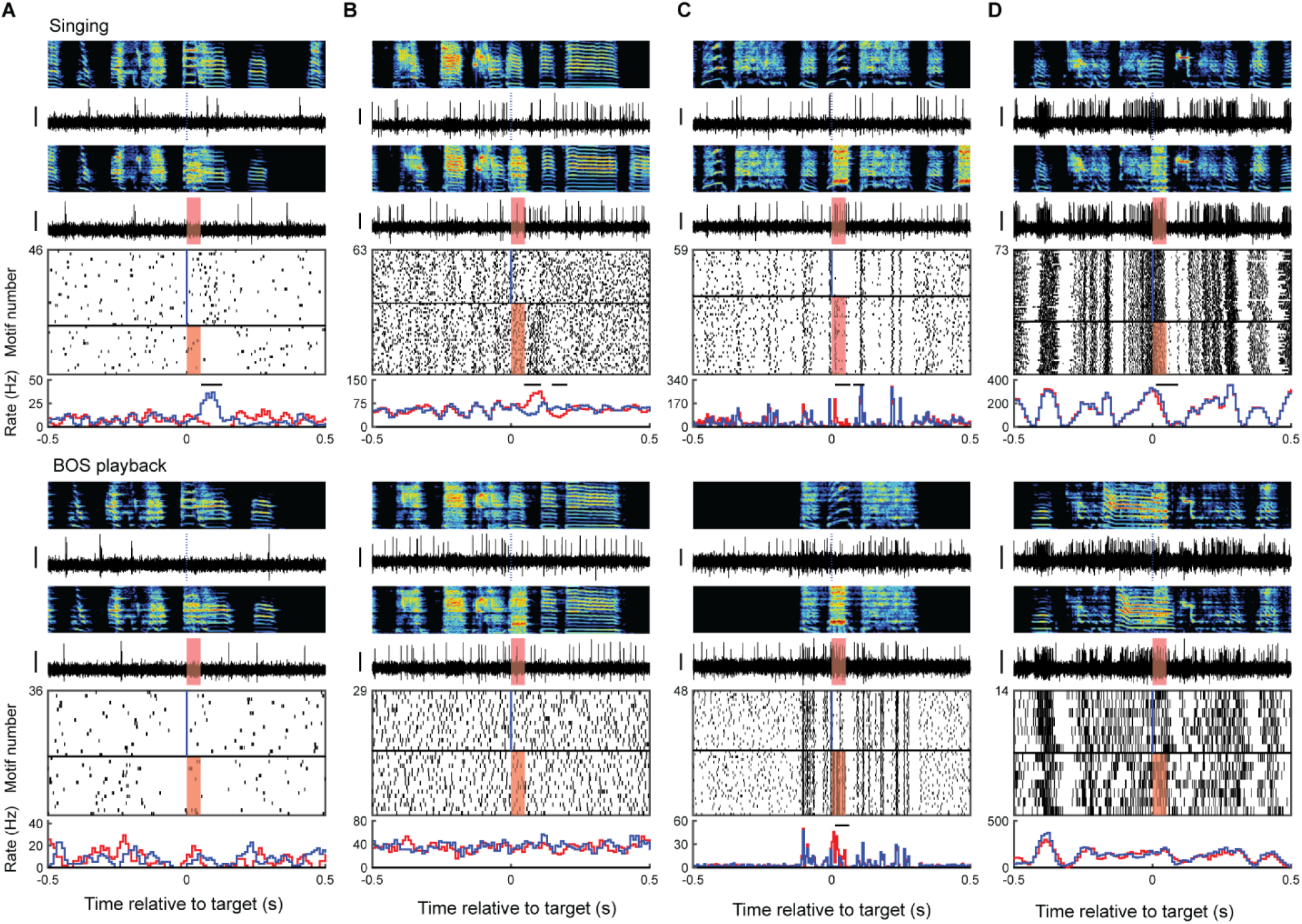
vBG responses during active singing and passive listening to bird’s own song. Examples of four neurons recorded during active singing, top row, and during passive playback of bird’s own song (BOS), bottom row. Data plotted as in Figure 3B for all neurons in both conditions. Shown are examples of neurons that exhibited error responses only during singing (A and B) and the only two putatively auditory neurons recorded in the dataset, which exhibited strong auditory responses to playback of BOS (C and D). Neuron in (B) is the vBGvta neuron from main Figure 5E.

**Figure S6.**
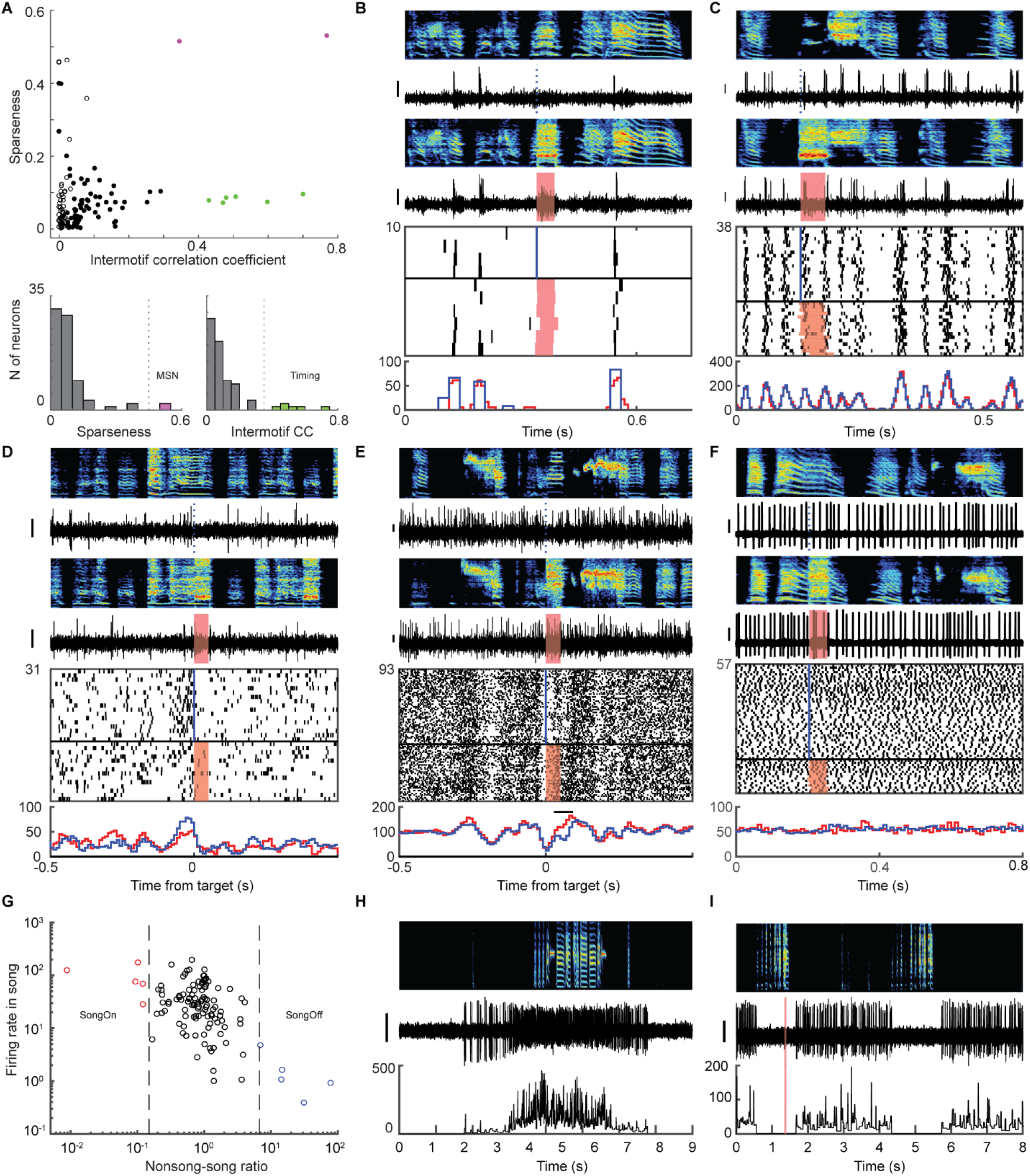
Examples of singing- and error-related representations in vBG. (A) Sparseness and intermotif correlation coefficient (ICC) distinguished two classes of time-locked vBG neurons. (B) MSN-like vBG neuron with sparse bursts time-locked to song motif. (C) vBG neuron with stereotyped high frequency bursts that tile the song motif. (D) Pre-target bursting neuron with high firing rate immediately before song target. (E) Pre-target pause neuron that was also error-activated. (F) vBG neuron unrelated to singing. (G) The ratio between mean firing rates outside and inside of singing identified vBG neurons gated by singing state. (H) Example song-activated neuron which fired at high rate during singing, but silent outside song. (I) Example song-suppressed neuron abruptly stopped firing during singing.

**Figure S7.**
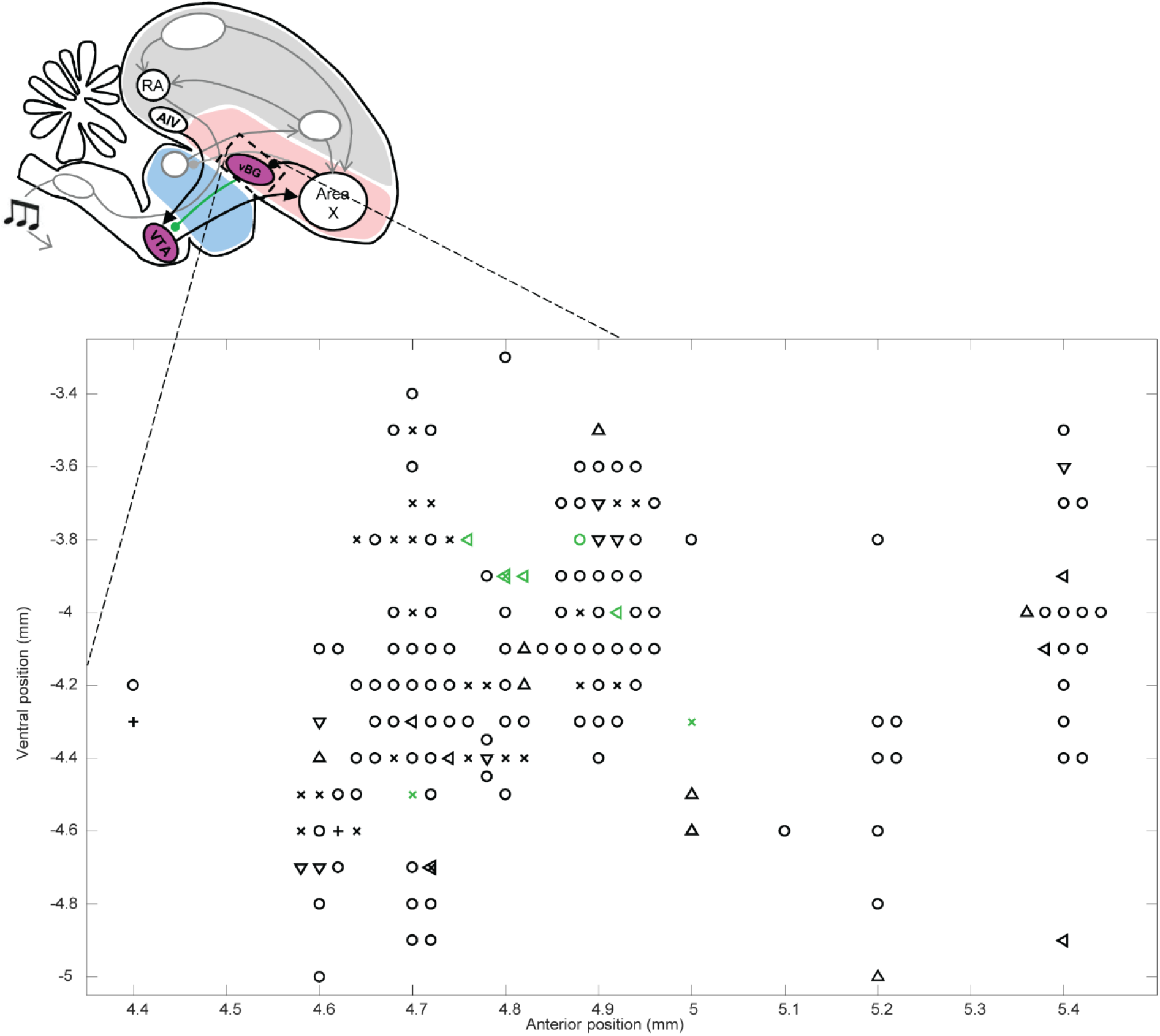
Diverse vBG cell types were spatially intermingled. Top, schematic of bird brain. Bottom, expanded view of the vBG region recorded, with the estimated position of each recorded neuron in the dataset. Mediolateral coordinates of 1.1 – 1.5 mm from midline are collapsed in this image. Cell type symbols: VTA-projecting (green), error responsive (x), MSN and burst-tiling (Δ), song-activated or song-suppressed (⍰), pre-target burst or pause (⍰), auditory (+), other (o).

